# nQuire: A Statistical Framework For Ploidy Estimation Using Next Generation Sequencing

**DOI:** 10.1101/143537

**Authors:** Clemens L. Weiß, Marina Pais, Liliana M. Cano, Sophien Kamoun, Hernán A. Burbano

## Abstract

Intraspecific variation in ploidy occurs in a wide range of species including pathogenic and nonpathogenic eukaryotes such as yeasts and oomycetes. Ploidy can be inferred indirectly - without measuring DNA content - from experiments using next-generation sequencing (NGS). We present nQuire, a statistical framework that distinguishes between diploids, triploids and tetraploids using NGS. The command-line tool models the distribution of base frequencies at variable sites using a Gaussian Mixture Model, and uses maximum likelihood to select the most plausible ploidy model. nQuire handles large genomes at high coverage efficiently and uses standard input file formats.

We demonstrate the utility of nQuire analyzing individual samples of the pathogenic oomycete *Phytophthora infestans* and the Baker’s yeast *Saccharomyces cerevisiae*. Using these organisms we show the dependence between reliability of the ploidy assignment and sequencing depth. Additionally, we employ normalized maximized log-likelihoods generated by nQuire to ascertain ploidy level in a population of samples with ploidy heterogeneity. Using these normalized values we cluster samples in three dimensions using multivariate Gaussian mixtures. The cluster assignments retrieved from a *S. cerevisiae* population recovered the true ploidy level in over 96% of samples. Finally, we show that nQuire can be used regionally to identify chromosomal aneuploidies.

nQuire provides a statistical framework to study organisms with intraspecific variation in ploidy. nQuire is likely to be useful in epidemiological studies of pathogens, artificial selection experiments, and for historical or ancient samples where intact nuclei are not preserved. It is implemented as a stand-alone Linux command line tool in the C programming language and is available at github.com/clwgg/nQuire under the MIT license.

## Introduction

Polyploidy, the presence of more than two complete sets of chromosomes, can under certain circumstances accelerate evolutionary adaptation by influencing the generation and maintenance of genetic diversity [1,2]. In addition, polyploidy also poses shortand long-term challenges to organismal fitness, which are associated with increased nuclear and cellular volume, propensity to aneuploidy, and disruption of gene expression regulation [3]. Interspecific comparisons between eukaryotic genomes can identify ancient polyploidization events. In contrast, more recent polyploidization events result in intraspecific variation of ploidy and, in some cases, aneuploidy. The presence of individuals of different ploidy in a population can hinder mating. Therefore, intraspecific ploidy variation tends to occur - although not exclusively - in organisms that have the capacity to reproduce asexually [4-6], are self-compatible or perennial [7].

Although ploidy traditionally has been investigated by measuring DNA content using flow cytometry [8], it can also be inferred from next generation sequencing (NGS) data either by examining k-mer distributions, or by assessing the distribution of allele frequencies at biallelic single nucleotide polymorphisms (SNPs) [4]. This methodology has been used to estimate ploidy in newly assembled genomes in order to identify the number of likely collapsed haplotypes on a per-contig basis [9], as well as to detect intraspecific variation of ploidy in the oomycete *Phytophthora infestans* [4,6] and in the Baker’s yeast *Saccharomyces cerevisiae* [5]. It also was successfully used for ploidy estimation in *P. infestans* historic herbaria samples that are not suitable for flow cytometry, since they do not contain intact nuclei [4]. The method assumes that alleles present at biallelic SNPs occur at different ratios for different ploidy levels, that is, 0.5/0.5 in diploids, 0.33/0.67 in triploids, and a mixture of 0.25/0.75 and 0.5/0.5 in tetraploids. To determine the ploidy level, the distribution of biallelic SNPs can be inspected visually [10], and/or qualitatively compared with simulated data [4]. However, this methodology does not provide summary statistics that permit quantifying how well the data fit the expected distributions, which is especially critical when dealing with noisy distributions typical for highly-repetitive genomes. An additional disadvantage of this approach is that it is preceded by the identification of variable sites (“SNP calling”), which is carried out using methodologies that benefit from a previously known ploidy level [11]. In a further development Gompert *et al.* [12] used biallelic SNPs in a Bayesian statistical approach to distinguish between ploidy levels from genotyping-by-sequencing data. This method was primarily developed for resequencing studies, where typically multiple individuals from populations with ploidy variation are genotyped, as it benefits from preexisting knowledge about the ploidy levels that may be observed. Based on the posterior probabilities emitted by the Bayesian model, this approach separates samples into ploidy clusters, using dimensionality reduction methods such as PCA. Since it allows the inclusion of training data of known ploidy, test samples can be assigned a ploidy level if they belong to a cluster that includes samples of known ploidy.

Here we present a statistical model that aims to distinguish between the distribution of base frequencies at variable sites for diploids, triploids and tetraploids, directly from read mappings to a reference genome. It models base frequencies as a Gaussian Mixture Model (GMM), and uses maximum likelihood to assess empirical data under the assumptions of diploidy, triploidy and tetraploidy. We evaluated the performance of our method for different sequencing coverages using published genomes of *S. cerevisiae* [5], and high-coverage genomes of *P. infestans* produced for this study.

## Methods

### Model and Implementation

We used base frequencies at variable sites with only two bases segregating to distinguish between diploids, triploids and tetraploids (Figure 1A). For that, we implemented a GMM that models the base frequency profiles as a mixture of three Gaussian distributions (Figure 1B), which are scaled relative to each other as:

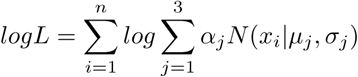

Here, *n* describes the numbers of data points, *x_i_* describes the value of each data point (i.e. the base frequency) and *μ_j_* and *σ_j_* are the parameters of the *j′th* of three Gaussian distributions *N_j_* that are scaled relative to each other through the parameter *α_j_*, with 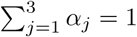.

**Figure 1.**
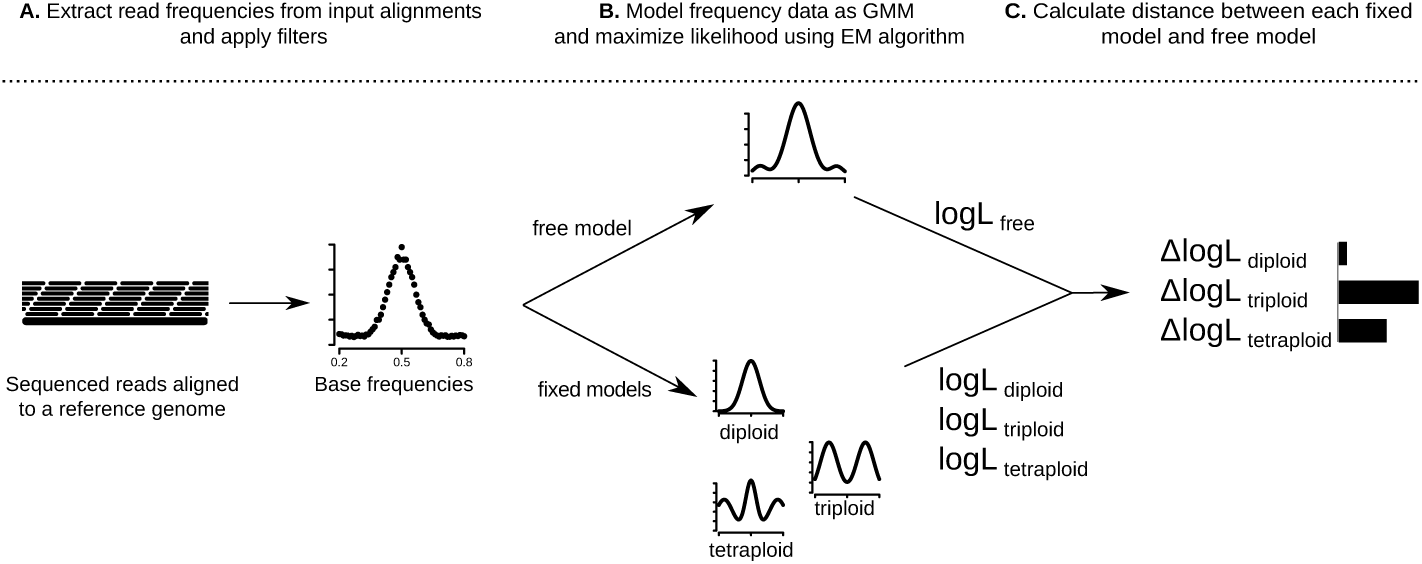
Overview of the Gaussian Mixture Model (GMM) based method used by nQuire to estimate ploidy. We illustrate the workflow using a diploid individual as an example. **A.** After sequenced reads are mapped to a reference genome, base frequencies are calculated at variable sites where only two bases are segregating. **B.** The base frequencies are modeled using a GMM and the likelihood is maximized using an Expectation-Maximization (EM) algorithm for both the free and the three fixed models (diploid, triploid and tetraploid). The maximized log-likelihoods (*logL*) are extracted for subsequent model comparison. The curves show a possible final state of the GMM under the assumptions of each of the four models. **C.** The Δ*logL* is calculated between the free model and each of the three fixed models (here represented as barplots). The fixed model with the smallest Δ*logL* is chosen as the true ploidy level (diploid in this example).

This model allows for estimating the parameters of the Gaussian mixture components, as well as their mixture proportions, by maximizing the log-likelihood, either with or without constraints on the possible parameter space.

The likelihood maximization of the GMM is implemented through an Expectation-Maximization (EM) algorithm (Figure 1B), which is specific to the GMM but can be extended to similar models. The algorithm estimates all parameters at once and computes a likelihood (“free model”). Alternatively, a likelihood can be calculated with constant parameters (”fixed models”) under the expectation of diploidy (one Gaussian with mean 0.5), triploidy (two Gaussians with means 0.33 and 0.67) and tetraploidy (three Gaussians with means 0.25, 0.5 and 0.75). Since all fixed models are nested within the free model, it is possible to directly compute the log-likelihood ratios, following:

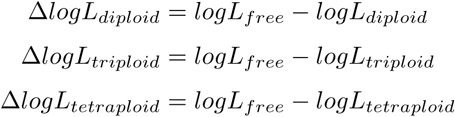

The Δ*logLs* describe the distance between each fixed model and the best fit under the assumptions of the GMM. A substantially lower Δ*logL* of one fixed model over the others supports the ploidy level described by this fixed model (Figure 1C). Therefore, we used Δ*logL* as summary statistics where the minimum value supports a given ploidy level.

Additionally, the GMM can be extended to a Gaussian Mixture Model with Uniform noise component (GMMU), by adding a uniform mixture component:

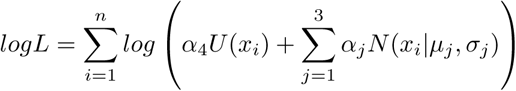

The constraint on the mixture proportions then becomes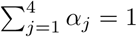.

The uniform noise component is used in our implementation to allow base-line noise removal. This is important when the Gaussian peaks are observable but embedded in a basal noise, which could be caused by highly repetitive genomes or low coverage.

### Multivariate Gaussian clustering

To cluster samples in three dimensions based on the normalized maximized log-likelihoods of the three fixed models, we used the mclust5 package [13] of the R programming language [14]. This package utilizes mixtures of multivariate Gaussian distributions to detect clusters in an arbitrary number of dimensions. mclust5 allows to set constraints on the volume, shape and orientation of each mixture component, by varying features of their covariance matrix either within each sample, or for all samples at once. For the analysis displayed in Figure 4, we used clusters of equal volume, but varying shape and orientation. This configuration represented the data the best, as assessed by the recovery of ploidy levels from cluster assignments.

### *Phytophthora infestans* libraries

The two benchmarking libraries from *P. infestans* were generated according to the protocol by Meyer and Kircher [15] from DNA extracted from lab cultures [16]. These libraries were sequenced to high coverage on an Illumina HiSeq 3000 machine in paired end 150bp mode. This sequencing data is available at the European Nucleotide Archive (ENA) under study number PRJEB20998.

## Results

### Performance

nQuire directly processes BAM files [17] and is designed to be efficient in memory usage and runtime. To process a 1GB *S. cerevisiae* BAM file (100x coverage), nQuire needs 70 seconds to build appropriate data structures, 6 seconds to run the models and calculate the maximum likelihood estimates, and uses a maximum of 8 Mb of RAM, whereas for processing a 10GB *P. infestans* BAM file (100x coverage) it needs 760 seconds, 100 seconds and 60 Mb of RAM, respectively.

### Analysis of individual samples

We evaluate nQuire’s performance using three *S. cerevisiae* samples at 100x coverage, which represent each of the three ploidy levels evaluated by the model, as well as two *P. infestans* samples, one diploid and one triploid, at 210x and 368x coverage, respectively. The Δ*logL* of each of the fixed models to the free model at full coverage is shown in Table 1. At those coverages, the Δ*logL* of the best model is more than two times closer to the free model than the second best. Additionally, it coincides in all samples with the ploidy level inferred by visually inspecting the empirical distributions of base frequencies at full coverage (Figure 2A-C and Figure 3A-B).

**Figure 2.**
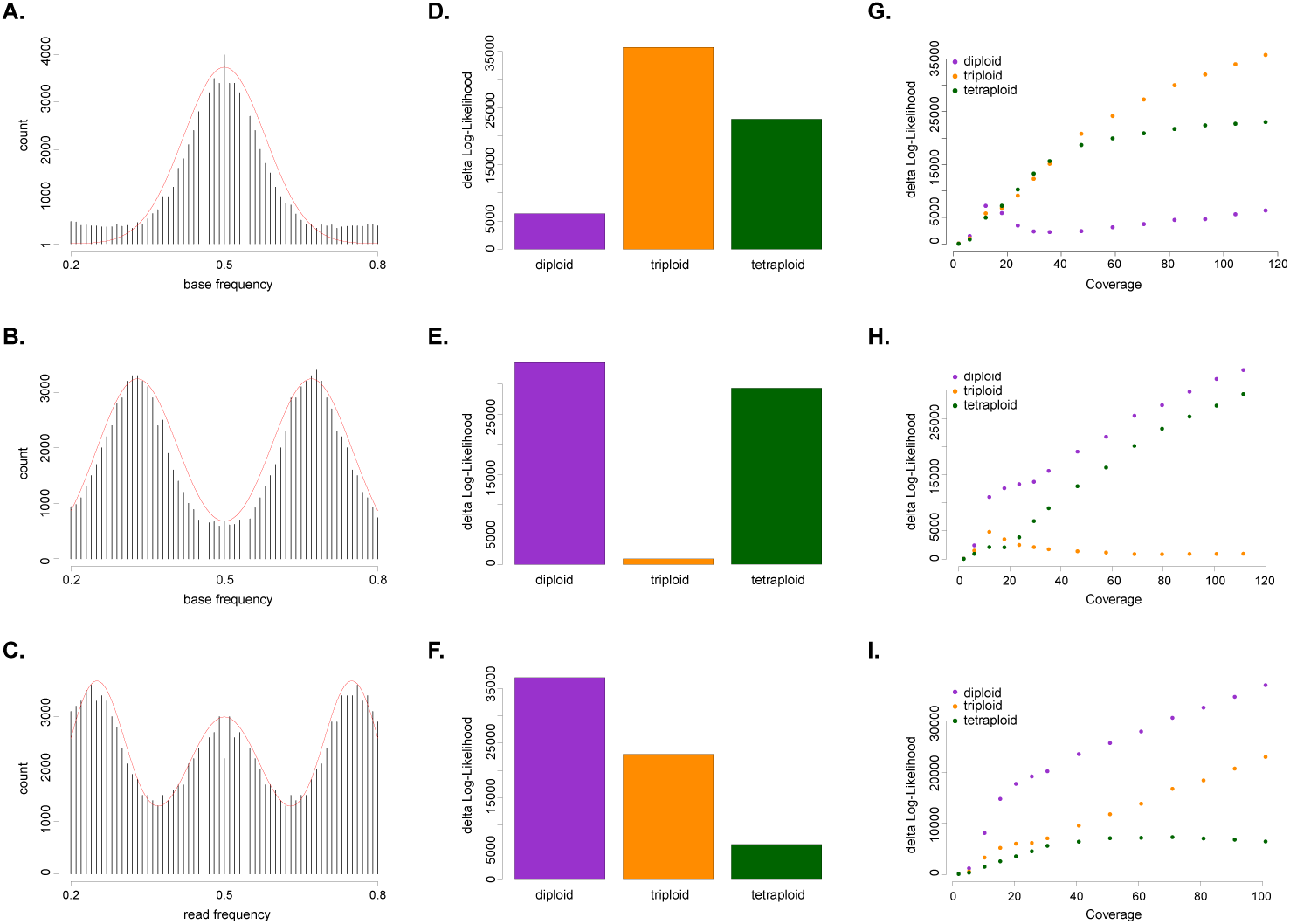
Evaluation and benchmarking of nQuire using *Saccharomyces cerevisiae*. Distribution of base frequencies at variable sites where only two bases are segregating for a diploid (**A.**), triploid (**B.**) and tetraploid (**C.**) sample. The barplots depict the Δ*logL* of all fixed models for the diploid (**D.**), triploid (**E.**) and tetraploid (**F.**) sample (also presented in Table 1). The plots depict the change of Δ*logL* of all fixed models as a function of genome coverage for the diploid (**G.**), triploid (**H.**) and tetraploid (**I.**) sample.

**Figure 3.**
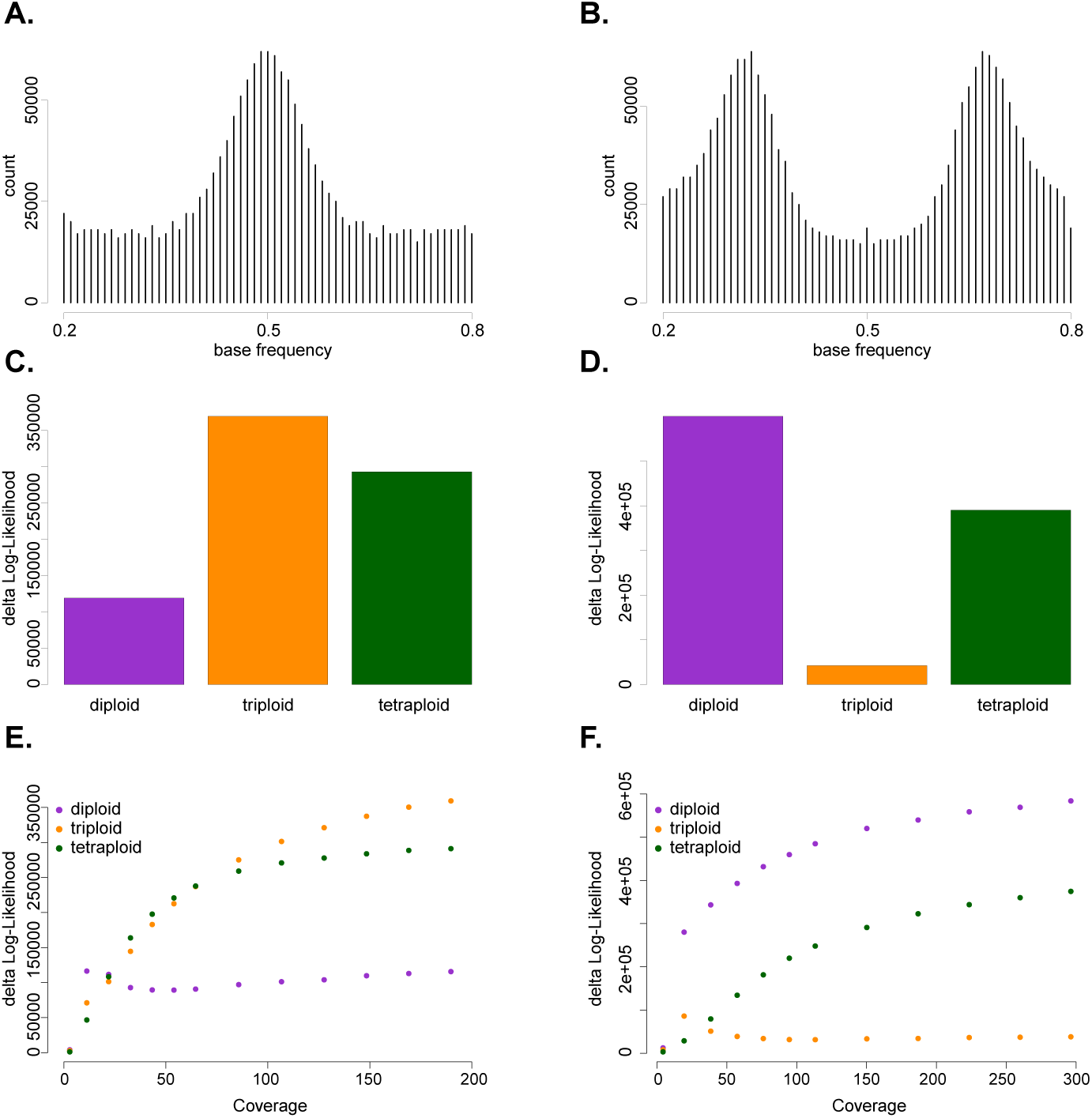
Evaluation and benchmarking of nQuire using *Phytophthora infestans.* Distribution of base frequencies at variable sites where only two bases are segregating for a diploid (**A.**) and a triploid (**B.**) sample. The barplots depict the Δ*logL* of all fixed models for the diploid (**C.**) and triploid (**D.**) sample (also presented in Table 1). The plots depict the change of Δ*logL* of all fixed models as a function of genome coverage for the diploid (**E.**) and triploid (**F.**) sample.

**Table 1.** Samples of *Saccharomyces cerevisiae* and *Phytophthora infestans* used to evaluate and benchmark nQuire. The smallest Δ*logL* for each sample is highlighted in bold.

| Sample | Ploidy | Species | Cov.<sup>+</sup> | $\Delta \log L_{2n}$ | $\Delta \log L_{3n}$ | $\Delta \log L_{4n}$ |
| --- | --- | --- | --- | --- | --- | --- |
| CBS7837 | 2n | *S. cerev.* | 116 | **6319** | 35721 | 23033 |
| CBS2919 | 3n | *S. cerev.* | 111 | 33614 | **920** | 29347 |
| CBS9564 | 4n | *S. cerev.* | 101 | 37003 | 22971 | **6429** |
| 99189 | 2n | *P. infes.* | 210 | **119218** | 369682 | 293194 |
| 88069 | 3n | *P. infes.* | 368 | 599933 | **42717** | 390002 |

<sup>+</sup> Average per-base coverage

### Coverage dependence

To investigate the impact of coverage on the performance of the GMM, we downsampled mapped reads from the *S. cerevisiae* (Figure 2G-I) and *P. infestans* (Figure 3E-F) strains shown in Table 1 to different coverage levels. These analyses showed that while the Δ*logLs* between the free model and the true fixed model start to plateau at low coverage, the Δ*logL* between the free model and the two incorrect fixed models keeps increasing at higher coverage (Figure 2G-I and Figure 3E-F).

### Analysis of population samples

In cases where multiple samples are sequenced simultaneously, it might be impractical to assess ploidy in each sample individually. In these cases, we propose to use maximized log-likelihoods of the three fixed models, normalized by that of the free model, to cluster samples into ploidy groups. The rationale is that within one species, the relative likelihoods of the fixed models will be similar within each ploidy level. As a proof of concept, we applied this to all di-, tri- and tetraploids from the *S. cerevisiae* test set [5], and clustered the samples into three groups in three dimensions using multivariate Gaussian clustering (see Methods). The sample set was manually scored for ploidy, and the overlap between clusters and manually assessed ploidy level was calculated (Figure 4). Running nQuire on raw data showed high recovery of ploidy level (93%, Figure 4A), which was further improved through our denoising implementation utilizing the GMMU (96%, Figure 4B).

**Figure 4.**
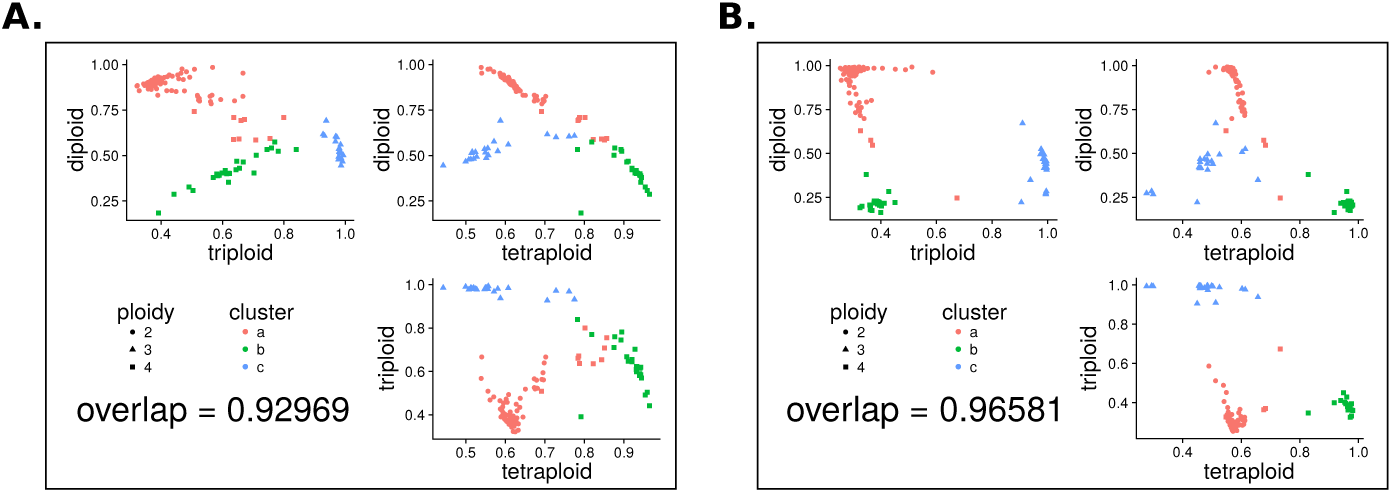
Clustering sets of samples into ploidy groups. Samples were clustered based on the normalized maximized log-likelihood of each fixed model representing each of the three ploidy levels. The clustering in three dimensions is shown with a set of three plots, one for each combination of ploidies. Shapes represent the manually assessed true ploidy levels, while colors show the assignments returned by the clustering algorithm. The agreement between the two is shown as a fraction. This analysis was conducted for samples before (**A.**) and after denoising (**B.**).

### Detection of Aneuploidies

Recent polyploidization is often associated with aneuploidies. To be able to detect those, nQuire allows to split the analysis of a sample by regions defined in BED format. We used this to reanalyze the sample YJM466 from the *S. cerevisiae* test set [5]. This sample had been shown to be triploid on whole genome level, but tetraploid for chromosome 6 and diploid for chromosome 9. The Δ*logLs* for the three fixed models individually calculated for each of the 16 chromosomes of *S. cerevisiae* confirmed this observation (Figure 5).

**Figure 5.**
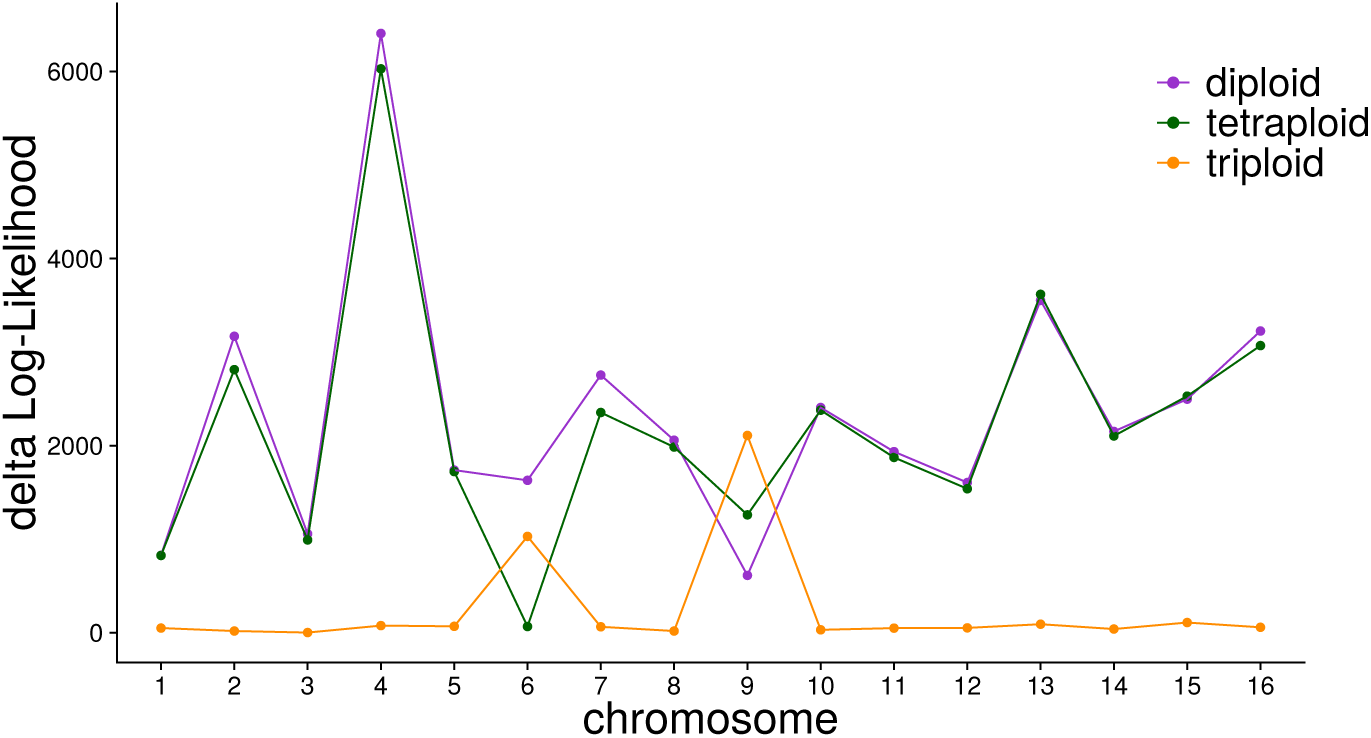
Detection of chromosome-wide aneuploidies. The ploidy estimation can be run for each chromosome independently, which enables detection of aneuploidies. The sample displayed here shows genome wide triploidy, but splitting the analysis by chromosome shows tetraploidy for chromosome 6 and diploidy for chromosome 9, as detected by Δ*logL*.

## Discussion

In addition to nucleotide and structural variation, certain organisms can also vary intraspecifically in their ploidy level, which constitutes another source of variation that selection might act upon. Using NGS data our method permits assessment of ploidy variation from data that is usually generated for variant detection. In contrast to previous methods that visually analyze the distributions of SNPs at biallelic heterozygous sites [4,6,10], we quantitatively distinguish between different ploidy levels based on the distribution of base frequencies at variable sites, using relative differences in likelihoods. In comparison to the approach proposed by Gompert and Mock [12], nQuire avoids the requirement of high quality SNP calls. The higher level of noise in the data resulting from this is accounted for by using Gaussian distributions. They approximate a binomial process, but are impacted less by the effects of outliers. Additionally, nQuire is a Linux command line tool that uses standard file formats as input and handles large genomes at high coverage efficiently.

In all test cases, triploids were the easiest to distinguish, most likely caused by the lack of probability density around 0.5 compared to the other two models. While diploids and tetraploids are more difficult to tease apart, our results on coverage dependence show that at sufficient coverage, the data fits the true model much better than either of the two alternatives (Figure 2G-I). For cases where the Gaussian peaks were largely overlapped by uniform noise, we extended our free model to include a uniform component, whose mixture proportion can be used - after likelihood maximization - for base-line removal. We show that this procedure improves the recovery of the true ploidy level when samples are clustered based on maximized likelihoods under the assumptions of the fixed models.

## Conclusion

We present nQuire, a statistical approach to distinguish diploids, triploids and tetraploids of recent evolutionary origin based on the distribution of base frequencies at variable sites. The method facilitates analysis of ploidy in single samples, and we demonstrate how to apply it to population scale data, when available. nQuire can also interact with BED files, to limit the analysis to certain sequence features, or divide it by regions of the genome, for example to detect aneuploidies. Our approach will be useful to assess intraspecific variation in ploidy from both historic and modern samples, as well as in experimental evolution experiments.

## Authors’ contributions

CLW, SK and HAB conceived the study. MP, LMC and CLW generated sequencing data. CLW developed the software application and analyzed the data. CLW and HAB wrote the the manuscript with input from all authors.

## Acknowledgments

We thank Michael Dannemann, Richard Neher, Thomas Mailund, Oliver Kohlbacher, Moises Exposito-Alonso, Kay Pruefer, and members of the Research Group for Ancient Genomics and Evolution (AGE) for useful discussions and input on model implementation; Patricia Lang, the AGE group and Michael Dannemann for comments on the manuscript; and the Presidential Innovation Fund of the Max Planck Society for financial support.

## References

1. A. M. Selmecki, Y. E. Maruvka, P. A. Richmond, M. Guillet, N. Shoresh, A. L. Sorenson, S. De, R. Kishony, F. Michor, R. Dowell, and D. Pellman, “Polyploidy can drive rapid adaptation in yeast,” Nature, vol. 519, pp. 349-352, Mar. 2015.

2. S. Venkataram, B. Dunn, Y. Li, A. Agarwala, J. Chang, E. R. Ebel, K. Geiler-Samerotte, L. Hérissant, J. R. Blundell, S. F. Levy, D. S. Fisher, G. Sherlock, and D. A. Petrov, “Development of a comprehensive Genotype-to-Fitness map of Adaptation-Driving mutations in yeast,” Cell, vol. 166, pp. 1585-1596.e22, Sept. 2016.

3. L. Comai, “The advantages and disadvantages of being polyploid,” Nat. Rev. Genet., vol. 6, pp. 836-846, Nov. 2005.

4. K. Yoshida, V. J. Schuenemann, L. M. Cano, M. Pais, B. Mishra, R. Sharma, C. Lanz, F. N. Martin, S. Kamoun, J. Krause, M. Thines, D. Weigel, and H. A. Burbano, “The rise and fall of the phytophthora infestans lineage that triggered the irish potato famine,” Elife, vol. 2, p. e00731, May 2013.

5. Y. O. Zhu, G. Sherlock, and D. A. Petrov, “Whole genome analysis of 132 clinical saccharomyces cerevisiae strains reveals extensive ploidy variation,” G3, vol. 6, pp. 2421-2434, Aug. 2016.

6. Y. Li, H. Shen, Q. Zhou, K. Qian, T. van der Lee, and S. Huang, “Changing ploidy as a strategy: The irish potato famine pathogen shifts ploidy in relation to its sexuality,” Mol. Plant. Microbe. Interact., vol. 30, pp. 45-52, Jan. 2017.

7. S. P. Otto and J. Whitton, “Polyploid incidence and evolution,” Annu. Rev. Genet., vol. 34, pp. 401-437, 2000.

8. S. Dirihan, P. Terho, M. Helander, and K. Saikkonen, “Efficient analysis of ploidy levels in plant evolutionary ecology,” Caryologia, vol. 66, pp. 251-256, Sept. 2013.

9. G. R. A. Margarido and D. Heckerman, “ConPADE: genome assembly ploidy estimation from next-generation sequencing data,” PLoS Comput. Biol., vol. 11, p. e1004229, Apr. 2015.

10. R. Augusto Corrêa Dos Santos, G. H. Goldman, and D. M. Riano-Pachon, “ploi-dyNGS: visually exploring ploidy with next generation sequencing data,” Bioinformatics, vol. 33, pp. 2575-2576, Aug. 2017.

11. G. A. Van der Auwera, M. O. Carneiro, C. Hartl, R. Poplin, G. Del Angel, A. Levy-Moonshine, T. Jordan, K. Shakir, D. Roazen, J. Thibault, E. Banks, K. V. Garimella, D. Altshuler, S. Gabriel, and M. A. DePristo, “From FastQ data to high confidence variant calls: the genome analysis toolkit best practices pipeline,” Curr. Protoc. Bioinformatics, vol. 43, pp. 11.10.1-33, 2013.

12. Z. Gompert and K. E. Mock, “Detection of individual ploidy levels with genotyping-by-sequencing (GBS) analysis,” Mol. Ecol. Resour., Feb. 2017.

13. L. Scrucca, M. Fop, T. B. Murphy, and A. E. Raftery, “mclust 5: Clustering, classification and density estimation using gaussian finite mixture models,” R J., vol. 8, pp. 289-317, Aug. 2016.

14. R Development Core Team, “R: A language and environment for statistical computing,” 2008.

15. M. Meyer and M. Kircher, “Illumina sequencing library preparation for highly multiplexed target capture and sequencing,” Cold Spring Harb. Protoc., vol. 2010, p. db.prot5448, June 2010.

16. D. E. L. Cooke, L. M. Cano, S. Raffaele, R. A. Bain, L. R. Cooke, G. J. Etherington, K. L. Deahl, R. A. Farrer, E. M. Gilroy, E. M. Goss, N. J. Grünwald, I. Hein, D. MacLean, J. W. McNicol, E. Randall, R. F. Oliva, M. A. Pel, D. S. Shaw, J. N. Squires, M. C. Taylor, V. G. A. A. Vleeshouwers, P. R. J. Birch, A. K. Lees, and S. Kamoun, “Genome analyses of an aggressive and invasive lineage of the irish potato famine pathogen,” PLoS Pathog., vol. 8, p. e1002940, Oct. 2012.

17. H. Li, B. Handsaker, A. Wysoker, T. Fennell, J. Ruan, N. Homer, G. Marth, G. Abecasis, R. Durbin, and 1000 Genome Project Data Processing Subgroup, “The sequence Alignment/Map format and SAMtools,” Bioinformatics, vol. 25, pp. 2078-2079, Aug. 2009.

